# Field and lab phenomics facilitate detection of genetic variation for iron deficiency chlorosis tolerance in sorghum

**DOI:** 10.64898/2026.04.01.715717

**Authors:** Gina Cerimele, Mitchell Kent, Michael Miller, Rex Best, Cleve Franks, Naqeebullah Kakar, Terry Felderhoff, Sarah Sexton-Bowser, Geoffrey P. Morris

**Affiliations:** Department of Soil & Crop Science, Colorado State University, Fort Collins, CO, USA 80523; Innovative Seed Solutions, Zionsville, IN, USA 46077; Department of Agronomy, Kansas State University, Manhattan, KS, USA 66506

## Abstract

Bioavailability of iron, an essential micronutrient to plants, is low in alkaline or calcareous soils, which are prevalent across semi-arid production regions. Breeding efforts to increase tolerance to iron deficiency chlorosis (IDC) in sorghum, a major crop of semi-arid regions, are confounded by spatial variation of stress severity in field trials. Here we developed and validated two high-throughput phenotyping approaches to address this challenge, with multi-spectral aerial imaging in the field and a controlled-environment assay to isolate the effects of iron bioavailability. In the field, severity and uniformity of stress are highly predictive of genetic signals for IDC tolerance (*R^2^* > 0.6 for soil pH metrics and *H*^2^). Plot-level data filtering for stress conditions based on control genotypes successfully addresses field spatial variation (unfiltered *H^2^* = 0.18 vs. filtered *H^2^* = 0.4). The controlled-environment assay proxies field stress using iron sources with differential bioavailability, evidenced by high heritability ( *H^2^* = 0.98) and phenotypic differential for hybrid control genotypes that matches field performance. Finally, we show that assay phenotypes are suitable for genome-wide association studies in global germplasm. Together, these field and lab phenomic approaches can be deployed to understand genetics of IDC tolerance and develop crops resilient to alkaline soils.

**HIGHLIGHT:** Stress severity and uniformity greatly impact detection of genetic signals underlying iron deficiency chlorosis tolerance in sorghum. A controlled-environment assay reduces spatial heterogeneity and improves assessment of tolerance genetics.

## INTRODUCTION

Iron is an essential micronutrient in plant nutrition, critical to chlorophyll synthesis, enzyme activation, cellular respiration, and other cellular processes (Briat et al., 2007; Connorton et al., 2017; Kobayashi & Nishizawa, 2012; Murgia et al., 2022). Iron deficiency inhibits photosynthetic productivity due to reduction of chlorophyll (Sharma, 2007), constraining crop growth and development and reducing yields (Barton et al., 2007). Iron deficiency-associated chlorophyll reduction in leaf tissue results in interveinal chlorosis, a characteristic yellowing between vascular elements (W. B. Anderson, 1982). Severe iron deficiency chlorosis (IDC) results in complete yellowing of leaf and shoot tissues, effectively halting photosynthetic productivity and, in turn, plant productivity (Mengel, 1994). While plant breeding and trait discovery for IDC tolerance achieved notable progress in crops like soybean (Aksoy et al., 2017; Lauter et al., 2020; Merry et al., 2019) and peanut (Pattanashetti et al., 2020; Tayade et al., 2022), relatively few other susceptible crops have comparable improvement in tolerance.

Soils are abundant in iron from parent material, however, plant iron bioavailability is dependent on other soil characteristics (Colombo et al., 2014). Calcareous soils are highly associated with incidence of IDC, where larger quantities of calcium carbonate (up to 15% of the soil) cause soil pH to rise and speciate iron from the bioavailable ferrous (Fe ^2+^) form to the unavailable ferric (Fe^3+^) form (Loeppert & Clarke, 1984). Iron bioavailability begins to decline as soil pH surpasses 6.0, resulting in nearly complete unavailability at a soil pH >7.5 (Lindsay & Schwab, 1982). To acquire ferric iron from the soil, plants evolved two main strategies that are induced by iron deficiency: a reduction-based Strategy I, and a chelation-based Strategy II (Kobayashi & Nishizawa, 2012; Marschner et al., 1986). Aside from grass species, all plants are believed to use Strategy I, where a plant secretes protons into the rhizosphere to lower the pH, speciating iron to the bioavailable ferrous form for ease of uptake. Grasses utilize Strategy II by synthesizing and exporting a family of metabolites called phytosiderophores, specifically mugineic acids (MA), to the rhizosphere where MAs bind ferric iron and the MA-Fe chelate complex is imported back into the root (Kawai et al., 1988; Marschner & Römheld, 1994). Efficiency of plant iron acquisition and utilization is a major driver of crop viability on high pH and calcareous soils.

Grain sorghum (*Sorghum bicolor*) is an economically and ecologically important crop in the semi-arid western and southern Great Plains region of the United States due to innate drought-tolerance characteristics (Monk et al., 2014) and ability to yield on stress-prone lands. However, sorghum performs poorly under iron deficiency (Figure 1A-B) when compared with other commodities such as maize or barley in a variety of metrics, including yield, biomass accumulation, and mugineic acid biosynthesis (Clark, 1982; Clark et al., 1988; Kawai et al., 1988; Onyezili & Ross, 1993). Breeding sorghums resilient to IDC is imperative to stabilize production acres in the western and southern Plains, where incidence of high pH and calcareous soils is common (Figure 1C) and may be further exacerbated by declining groundwater (Clark, 1982; Hirmas & Mandel, 2017). Though the genetic basis of IDC tolerance in sorghum has yet to be characterized, the homology of cloned iron homeostasis genes from other species can be leveraged to identify candidate breeding targets (Figure 1D) (Aung et al., 2019; Bashir & Nishizawa, 2006; Curie et al., 2001; Higuchi et al., 1999; Kobayashi & Nishizawa, 2012; Nakanishi et al., 1993; Nozoye et al., 2011, 2015, 2019; Ogo et al., 2007, 2008; Suzuki et al., 2006; Takahashi et al., 1999, 2001; Wang et al., 2020). Phenotypic selection for IDC tolerance in sorghum breeding nurseries is not reliable due to the spatial variability of IDC-associated factors in the soil landscape causing unequal plant stress (Bloom & Inskeep, 1986; D. R. Morris et al., 1990). Therefore, breeders cannot be sure that a plot is performing well due to adaptive genetics or simply due to minimal stress (Williams et al., 1987). Thus, there is a demand for a controlled-environment method to screen sorghum breeding material for IDC tolerance under uniform stress which proxies the field environment.

**Figure 1.**
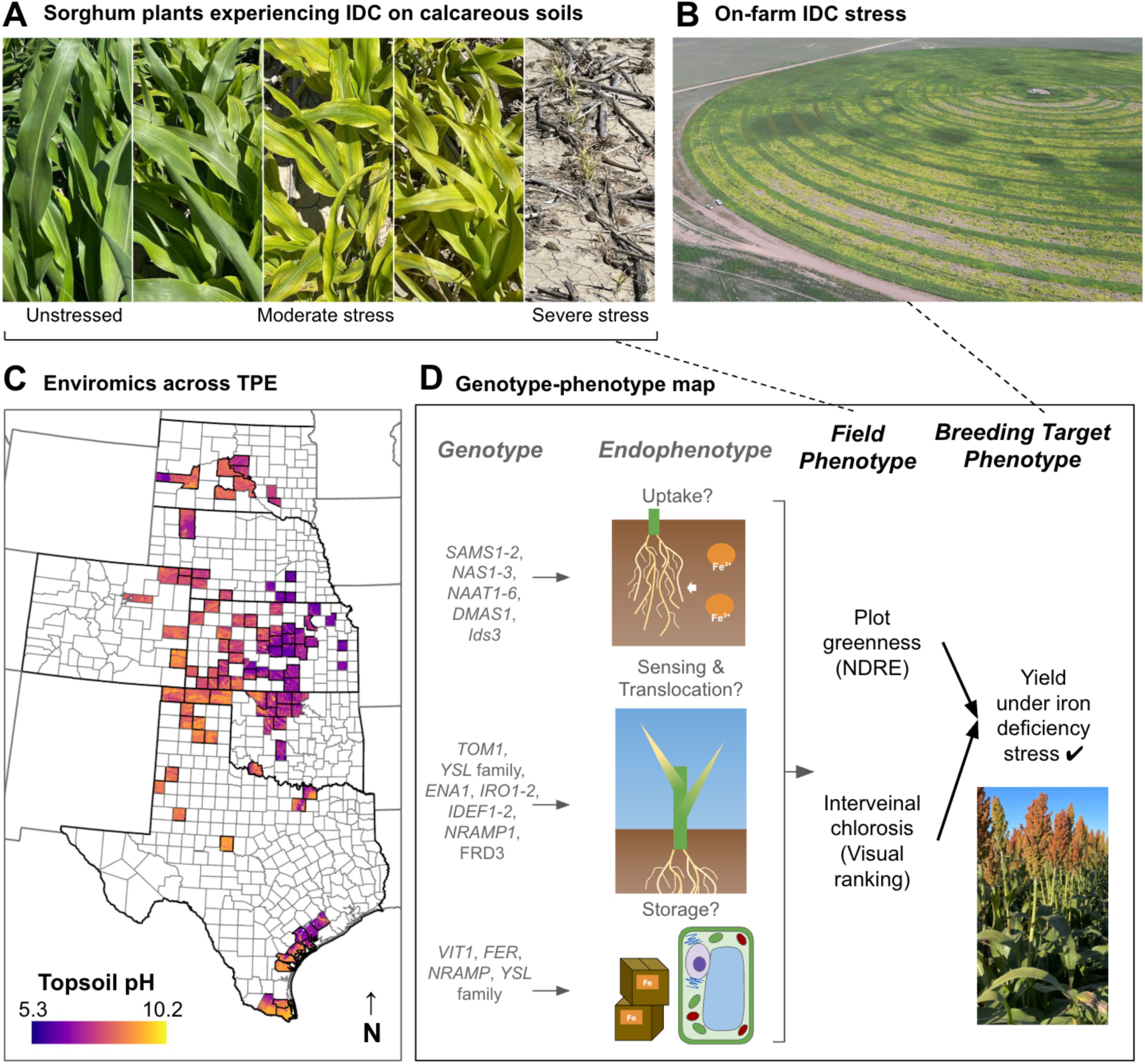
Sorghum improvement for tolerance to Iron deficiency chlorosis is necessary to stabilize cropping viability across the target market area. **A)** The scale of severity of iron deficiency chlorosis symptoms in sorghum under field conditions. All photos were captured on the same day from sorghum genotypes showing variable IDC tolerance. **B)** Sorghum hybrid seed production field severely impacted by iron deficiency where the green bands are planted with an IDC tolerant male pollen parent and the yellow or barren soil were planted with an IDC susceptible female seed parent. **C)** Soil pH, a soil property strongly associated with incidence of iron chlorosis, across the target population of environments (TPE) for grain sorghum breeding programs in the Sorghum Belt of the United States. Counties with any color represent counties with sorghum acreage planted in 2024 according to the United States Department of Agriculture National Agricultural Statistics Service. **D)** The hypothesized genotype to phenotype map emphasizing genes important to iron homeostasis pathways in grasses and their application towards phenomics benefiting the genetic improvement of iron deficiency chlorosis tolerance in sorghum. Numbers in gene names separated by a hyphen (e.g. *NAS1-3*) indicate sequentially named members of a gene family (e.g. *NAS1*, *NAS2*, *NAS3*).

Phenomic tools and analyses can bridge the gap between spatially-confounded field experiments and reductionist lab approaches (York, 2019). Field experiments assess on-farm performance across the target population of environments (TPE), while lab experiments isolate specific stresses to unveil biological mechanisms while still applying environmental relevance. To proxy the field environment for IDC tolerance phenotypic segregation, iron sources in sufficient and deficient treatments should include ferrous (Goos & Germain, 2001) and ferric (Olsen, 1958) iron sources, respectively. Iron deficiency assays used in molecular genetics studies commonly formulate deficient treatments by removing iron from the system, or reducing the amount of ferrous iron applied relative to a sufficient treatment (Chan-Rodriguez & Walker, 2018; Higuchi et al., 2024; Nakib et al., 2021; Prity et al., 2021; Sharma, 2007). While useful to dissect molecular mechanisms of response to iron starvation and deficiency, these methods do not proxy field relevant evaluation of germplasm performance for IDC tolerance based on genetic variation driving the ability of the plant to acquire biolimiting ferric iron from the rhizosphere.

In this study, we developed a controlled-environment assay that aids breeding decisions for sorghum IDC tolerance by screening plant performance under field-relevant, low bioavailability iron stress. We found the assay improved detection of genetic signals underlying IDC tolerance relative to field experiments, and scaled to screen elite or exotic germplasm for candidate IDC tolerance trait donors. The assay design elements satisfy a high-throughput and reproducible method to evaluate breeding germplasm for increased tolerance; unlocking a key bottleneck in developing IDC-tolerant sorghums. Integration of controlled-environment and lab screening methods with on-farm field assessment supports crop improvement efforts for increased IDC tolerance in sorghum.

## MATERIALS AND METHODS

### Field data collection

An irrigated field experiment to assess genotype performance under IDC stress was planted in summer 2025 in Tribune, KS due to historic severe stress pressure at this site. Field configuration included 62 ranges by 64 columns comprising 4.25 m long x 0.75 m wide, single row plots where test genotypes were randomized and planted between alternating columns of positive and negative control hybrid plots (Figure 2A) used to calibrate spatial stress severity. A single positive control hybrid (putative tolerant to IDC stress) and single negative control hybrid (putatively susceptible to IDC stress) were used in replication throughout the entire field. Field conditions were maintained as well-watered throughout the growing season.

**Figure 2.**
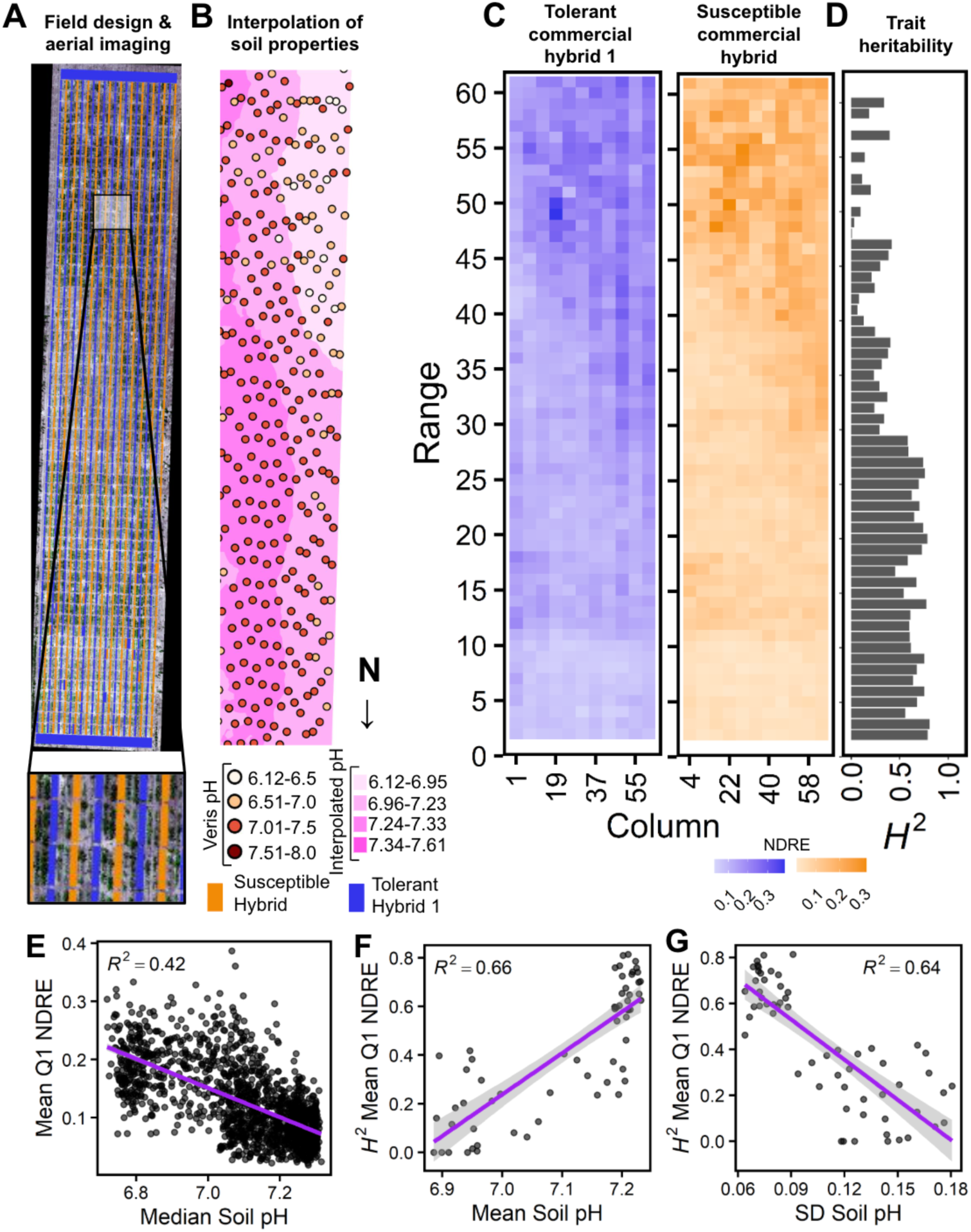
Soil spatial variability inhibits the ability to assess sorghum resilience to iron deficiency chlorosis in managed field screening. **A)** A sorghum breeding program IDC tolerance screening nursery utilizes strip-planted tolerant and susceptible control hybrids spanning the length of the field to calibrate nearby plot stress severity for test entries. Field layout is formatted T-test-test-S-test-test-T-test-test-S… where ‘T’ is the tolerant control hybrid 1, ‘S’ is the susceptible control, and ‘test’ is any test entry under evaluation. **B)** High-throughput proximate soil sensing using a tractor-drawn Veris sensor system provides georeferenced point data of soil pH measurements, facilitating interpolation of soil pH across the field landscape. **C)** Putatively tolerant (blue) and susceptible (orange) commercialized control hybrids with stable genetics show variability in iron deficiency chlorosis sensitivity phenotype proxy, normalized difference red edge (NDRE), in field experiments. Each square represents a single field plot unit, where a color family contains only entries from a single control hybrid to compare within-genotype phenotypic variation. Test entries and the alternate control were omitted for each visualization. Darker squares indicating higher NDRE suggest ambiguous interpretation as more tolerant to iron deficiency stress OR less iron deficiency stress impacting that plot. **D)** Distribution of broad-sense heritabilities (*H*^2^) for control hybrids in each range of the field. **E)** The correlation of plot-level soil median pH data with plot-level NDRE IDC tolerance phenotypes indicates a strong negative relationship between rising pH and plant performance. **F)** Strong positive correlation between field range-level means of plot-level median soil pH and the corresponding rangle-level broad-sense heritability estimates of NDRE suggest that rising pH strengthens detection of genetic signals for IDC tolerance. **G)** Strong negative correlation between field range-level standard deviation in plot-level median soil pH and range-level broad-sense heritability estimates of NDRE suggests that increasing variability in soil pH weakens detection of genetic signals for IDC tolerance.

A tractor-drawn Veris MSP3 soil mapping system (Veris Technologies, Kansas, USA) was used to collect soil pH measurements on the field (22 May 2025) prior to spring planting. The pH point data were used to generate an interpolated soil pH raster of the field via ordinary kriging (Liu et al., 2006; McBRATNEY & Webster, 1983) in ArcGIS Pro (Esri Inc., California, USA). A plot shapefile specifying the geographic boundaries of each field plot was used to extract zonal statistics on the interpolated soil pH data for each control plot. A DJI M300 drone and Sentera 6X sensor were used to phenotype the field for IDC tolerance 36 days after planting.

Orthomosaics were stitched using structure from motion software Agisoft Metashape (Agisoft, 2026). Soil pixels were removed from the plots using an Excess Green (ExG) index (Meyer & Neto, 2008) vegetation filter where pixels < 0.03 ExG were converted to no data, resulting in retention of only vegetation pixels in subsequent phenotype extraction. A normalized difference red edge (NDRE) index was then applied, where the mean of the first quartile (Q1) of NDRE values for each plot was extracted as the IDC stress phenotype to capture the portion of the plot that experienced the most severe stress response (Williams et al., 1987). The NDRE index was chosen due to its increased sensitivity to chlorophyll content over NDVI (Torino et al., 2014). A higher NDRE value indicates a greener plant. In addition to use of premergent herbicide, the field was hand-weeded before aerial phenotype collection to minimize noise from weed vegetation pixels.

### Field data analysis

R statistical software (v4.4.3, R Core Team 2025) was used for all statistical analyses across the study, with the lme4 R package (Bates et al., 2015) used for all linear models. The broad-sense heritability (*H^2^*) (Equation 1) of the mean Q1 NDRE was calculated for the control plots in each range spanning the field. Ranges with low and high *H^2^* were assumed to indicate low and high IDC stress pressures, respectively. A linear regression and plot-level correlation between the median soil pH of the plot from the extracted soil zonal statistics and NDRE phenotype was completed. A linear regression was also performed for the range-level correlation between *H^2^*and mean of the median soil pH for the plots in the range. For each range the *H^2^* of mean Q1 NDRE was calculated along with the mean of the median soil pH of plots in the range. Finally, a simple linear regression and range-level correlation between *H^2^* and standard deviation of the median soil pH for the plots in the range was completed. For each range, the *H^2^*of mean Q1 NDRE was calculated along with the standard deviation of the median soil pH of plots in the range.

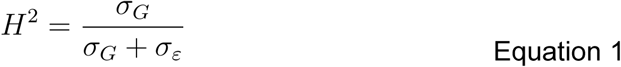

Phenotypic distributions of mean Q1 NDRE for control hybrids were assessed in the field for three methods of plot stress filtering. The first method used retention of all plots in the field (no filter). The second method (‘neighbor-filtering’) leverages the alternating strip-planted controls where a pair of control plots are removed if, within the pair, the susceptible control is performing equal to or better than the tolerant control. The third method (‘pH filtering’) removed any plot where the interpolated plot-level soil median is lower than an induced stress threshold of 7.1. *H^2^* of NDRE was calculated for control hybrids after filtering in all methods.

### Set-up and operation of controlled-environment IDC assay

Individual plants are isolated to 50 mL centrifuge tubes to prevent sharing of mugineic acids. A small hole is drilled in the bottom of each tube to provide drainage and mitigate molding. Three medium-sized cotton balls are then placed loosely into the bottom of each tube to provide substrate for root growth and nutrient solution retention. Cotton is used as a growth substrate to limit potential introduction of trace exogenous iron that is commonly found in substrates like sand, calcined clay (e.g. Turface), and commercial potting mixes. Two seeds of the test genotype are sandwiched between an additional two cotton balls and placed into the tube with the “seam” of the two cotton balls facing up to facilitate easier germination and root penetration through the cotton. Two seeds are used to increase the chance of a germinated sample. The tubes are then placed into the cells of 128-cell nursery plug trays to remain stable and upright during the growth period, and placed onto a standard greenhouse bench.

After placing the tubes into the plug trays, DI water is used to saturate the cotton in each tube to stimulate germination. The tubes are then loosely covered with heavy paper to darken the environment during germination. At day 8 following initial saturation when seeds have germinated, application of nutrient treatments begins. A 10mL pipette is used to apply 10mL of the designated iron treatment nutrient solution to each tube in the treatment class. Subsequent treatment applications occur every three days until phenotyping. If cotyledons or leaves are stuck in the cotton, forceps are used to free them. Any tubes with no germination are removed after two weeks.

### Nutrient formulations for controlled-environment IDC assay

A base nutrient solution is created to supply all essential macro- and micronutrients to plants in sufficient quantities, other than iron which is added later to specify a sufficient or deficient treatment. The base nutrient solution is prepared based on nutrition guidelines for sorghum and formulations adapted from other similar iron deficiency experiments. The base solution consisted of the following nutrients and concentrations: 4mM Ca(NO _3_)_2_, 2mM MgSO_4_, 1.33mM (NH_4_)_2_HPO_4_, 1.25mM KNO_3_, 100µM NaCl, 10µM MnSO_4_, 1µM CuSO_4_, 1µM ZnSO_4_, 33µM H_3_BO_3_, 0.2µM Na_2_MoO_4_, and 0.1µM CoSO_4_. All iron sources are omitted from the base nutrient solution and DI water is used in the solution. Large quantities of the base solution are mixed and then divided equally to which different iron sources are then added to form the respective sufficient and deficient iron treatments. To form the sufficient treatment, Fe(III)-EDTA and FeSO _4_ are added to the subset base nutrient solution to achieve concentrations of 0.15mM and 0.1mM, respectively. To form the deficient treatment, Fe _2_(SO_4_)_3_ is added to the subset base nutrient solution to achieve a concentration of 0.15. Solution pH is adjusted for each treatment, where sufficient treatment pH is adjusted to 6.5-6.7 using HCl and deficient treatment pH is adjusted to 7.8-8.1.

### Greenhouse conditions for controlled-environment IDC assay

Greenhouse conditions are maintained at standard acceptable conditions for sorghum growth. Supplemental lighting via 400-watt high pressure sodium lights is applied to create a day and night cycle of 16 hours of day and 8 hours of night. Air temperature is maintained at 26-27℃ and 22-23℃ for daytime and nighttime temperatures, respectively. Humidity is maintained at ambient levels, approximately 25-30% relative humidity.

### Phenotyping for controlled-environment IDC assay

Samples are phenotyped at 28-31 days post-planting at the 3-leaf stage. Each sample is phenotyped for IDC using a quantitative relative chlorophyll content (SPAD) measurement (Peterson & Onken, 1992). SPAD phenotypes are collected using a SPAD 502 Chlorophyll Meter handheld unit (Spectrum Technologies, Inc.) where a higher value indicates greener leaf tissue. Six SPAD measurements are taken for each sample to account for some natural variation in tissue greenness and stress response; three measurements encompassing the base, middle, and end for each of the two youngest leaves that are able to be measured with the SPAD meter. SPAD measurements are then averaged to provide a single SPAD datapoint for each sample.

### Validation of controlled-environment assay as a field proxy

The tolerant and susceptible hybrid controls were assessed for performance under assay conditions. Due to changes in commercialization demand, the tolerant hybrid was updated to a different hybrid during the project, resulting in evaluation of two tolerant hybrids over the course of assay development (designated as ‘Tolerant Hybrid 1’ and ‘Tolerant Hybrid 2’). The assay was prepared and completed as described in the sections above. Phenotype values of hybrid control performance under sufficient and deficient treatments were visualized. *H^2^* of SPAD was calculated for hybrid control lines under deficient assay treatment and compared with *H^2^* for NDRE under field conditions.

### Deployment of controlled-environment IDC assay in global germplasm

A diversity panel of 330 georeferenced global accessions was grown in three replicates. The panel of georeferenced accessions maximizes spatial distribution, offering a diverse pool of genetic variation that may reveal genetic factors shaped by evolutionary processes across spatial gradients important to iron presence and bioavailability (G. P. Morris et al., 2026). Each replicate included all genotypes in two treatments: iron sufficient and iron deficient. Due to the volume of samples and throughput of the phenotype collection method, replicates were grown one after another at different timepoints as a complete block. SPAD measurements of the hybrid controls also grown during the experiment were plotted with the distribution of mean SPAD values averaged over the replicates for each test accession.

### Genome-wide association study of controlled-environment phenotypes

SPAD phenotypes from the controlled-environment assay screening of the georeferenced global accessions described above were utilized in conjunction with high-density single nucleotide polymorphism (SNP) genotype data for association mapping of IDC tolerance. Association analyses were only completed for the deficient treatment phenotypes. The phenotype input consisted of best-linear unbiased predictors (BLUPs) estimated for average SPAD using a linear model (Equation 2), where *y_ij_* is the SPAD value for genotype *i* in replication *j*, *g_i_* is the genotype random effect, *r_j_* is the random effect of replication, and *ε_ij_* is the residual error.

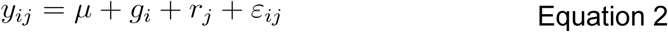

The genotype random effect was extracted as the BLUP. BLUPs were used due to the presence of unbalanced reps driven by variability in germination. A total of 312 unique accessions with sufficient phenotype data were considered in the analysis. Genotype data consisted of a resequencing SNP dataset. PLINK was used to complete linkage disequilibrium (LD) pruning with a SNP window of 50, step size of 5, and an *r^2^* threshold of 0.2. LD pruning reduced the dataset from 35,877,217 SNP markers to 9,769,987 SNP markers. GEMMA was used to calculate a relatedness matrix and run a univariate linear mixed model (LMM) association model. This analysis utilizes a centered relatedness matrix where it is not assumed that the effect size of a SNP depends on its minor allele frequency. GEMMA uses the relatedness matrix to fit a kinship (K) term in the LMM association model. A Bonferroni-adjusted threshold was calculated and used to evaluate meaningful association significance.

The genomic positions of sorghum homologs of *a priori* IDC tolerance candidate genes were obtained from the *Sorghum bicolor* v5.1 reference genome using Phytozome v13 (Goodstein et al., 2012). Positions of candidate genes were overlaid onto the Manhattan plots of association analysis results. Using the JBrowse functionality of Phytozome, any candidate genes visualized near distinct marker trait associations (MTA) were investigated for physical proximity of the MTA SNP to the candidate gene locus. Additionally, the five highest MTA SNPs on each chromosome were investigated for proximity to any gene, regardless of *a priori* candidate status.

## RESULTS

### Spatial variation for soil pH exists in managed field experiments

To assess the extent of spatial variation for relevant soil factors in a managed stress sorghum IDC tolerance breeding nursery, soil pH data interpolated from high-throughput proximate soil sensing data was evaluated across the field landscape. After generating an interpolated soil pH raster dataset (Figure 2B) from georeferenced field sampling points (Figure 2A) followed by extraction of plot-level statistics for all control hybrid plots, median soil pH for each plot was visualized according to the range and column orientation of the field layout (Figure 2C). The field pH ranged from 6.72 to 7.31 where plots in the southern section, and especially southwestern corner, of the field were characterized by a substantially lower median pH, ranging between 6.7 and 6.9. The median pH observed in plots along the entire eastern length of the field, and particularly in the northeastern section, maintained near the maximum pH of 7.3. The observed range of soil pH across the field supports our hypothesis that spatial variation for soil factors associated with IDC exists within our field site.

### Genetic differences in IDC tolerance are revealed when soil pH is high and uniform

To determine if differences in soil pH are contributing to phenotypic variation for IDC tolerance we assessed the relationship between plot-level median soil pH and the IDC tolerance phenotype, mean normalized difference red edge (NDRE) of the first quartile of vegetation pixels in a plot. It is expected that, given the speciation of ferric iron under pH of 7 and above, high pH would correlate with low NDRE, a proxy for plant vegetative growth. At the plot level, the *r* and *R*^2^ of median soil pH versus NDRE phenotype were -0.65 and 0.42, respectively, indicating nearly half of the variation in phenotype can be explained by soil pH (Figure 2E). Thus, while soil pH is somewhat predictive of plant performance, additional genetic and environmental factors may be playing important roles in sorghum IDC tolerance phenotypes.

To assess the relationship between spatial distribution of soil pH and consistency of genetic signals for IDC tolerance, the predictiveness of range-level soil statistics for broad sense heritability (*H^2^*) was determined. It is expected, given the speciation to ferric iron at pH 7 and above, that higher mean pH for a range would result in a higher *H^2^* for an IDC tolerance phenotype due to stress induction strengthening genetic signals. Similarly, a higher standard deviation for range-level pH would be expected to correlate negatively with *H*^2^ due to stress variability in a range masking genetic signals. First, we assessed the heterogeneity of *H^2^*for NDRE in the control hybrids across the ranges in the field. We found the range of *H^2^* spanned from zero to 0.81, indicating notable spatial variability in detection of genetic signals for IDC tolerance (Figure 2D). Next, the relationships between soil characteristics and genetic variation were assessed. At the range level, the *r* and *R*^2^ of mean soil pH versus *H^2^*of NDRE were 0.81 and 0.66 (Figure 2F), respectively, indicating that increased IDC stress severity with rising pH enhances detection of genetic signals for IDC tolerance. Furthermore, the *r* and *R*^2^ of the standard deviation of soil pH versus *H^2^* of NDRE were -0.8 and 0.64 (Figure 2G), respectively, demonstrating that greater spatial variability in IDC-associated soil characteristics weakens detection of genetic signals for IDC tolerance.

### New high-throughput controlled-environment assay isolates effects of iron bioavailability

To address the spatial challenges of field phenotyping, we developed a controlled-environment assay to isolate the effects of iron bioavailability on plant performance. The assay controls for these factors in several ways. Genotypes are isolated in their growth environment to prevent rescue of deficient phenotypes through mugineic acid sharing (Senoura et al., 2024). As a growth substrate, cotton is used to limit introduction of extraneous iron that is present in commercially available, mineral-based media. Two seeds are planted per sample and later thinned to one plant to control for genotypes with poor germination. A base nutrient solution containing all essential components of nutrition other than iron is prepared with distilled water to exclude exogenous iron then split into sufficient and deficient treatments where respective iron sources are added to limit batch effects related to off-target nutrients. The sufficient and deficient treatments utilize available Fe(III)-EDTA and FeSO _4_ and unavailable Fe_2_(SO_4_)_3_ iron sources, respectively, to not exclude genetic variation in iron uptake of ferric iron driving phenotypic response. Furthermore, pH is adjusted between the treatments to further proxy the field IDC stress conditions characteristic of calcareous soils. Averaging six relative chlorophyll content (SPAD) measurements across two leaves for each sample accounts for natural patterns in leaf greenness and stress symptoms.

To determine the cost effectiveness of the controlled-environment assay, a price per sample and initial startup costs were assessed alongside effort and timing for assay use. The main consumable materials, including centrifuge tubes, nutrient solution chemicals, cotton balls, and sample labels, amount to approximately $0.20 per sample. While highly dependent on the equipment available in a given lab or industry setting and the specific use plan for the assay, approximately $345 of initial start-up materials such as a basic pH electrode, basic 10mL pipette, containers for stock and bulk solutions, and other standard laboratory tools may be needed. It is reasonable to expect that, for one person working alone, approximately 30 hours would be needed for assay preparation for 500 samples. After plant germination, nutrient treatment application for 500 samples would require approximately 4-6 hours every third day. While the labor input is not insignificant, the flexibility to utilize the assay in a greenhouse setting any time of year facilitates important IDC tolerance insights outside of outdoor growing season constraints when labor may be more available for alternative activities.

### Consistent differentiation of tolerant vs. susceptible genotypes across field and lab approaches

To evaluate the newly developed assay as an alternative method to field screening for spatially-controlled phenotypic evaluation of IDC tolerance, we assessed the hybrid controls for phenotypic response to sufficient and deficient treatments where only the bioavailability of the source of iron and nutrient solution pH differed. Under sufficient iron treatment, we expected the hybrid controls to exhibit no IDC stress and perform similarly to one another. By contrast, under deficient iron treatment, it is expected that the susceptible hybrid performs significantly worse than the tolerant hybrids, which are expected to perform closely to the observed sufficient treatment phenotypes. The patterns observed in the controlled-environment assay mirror these expectations (Figure 4A), where the difference in greenness (SPAD) between the sufficient and deficient treatment means for the tolerant hybrids was 3.2 for tolerant hybrid 1 and 4.5 for tolerant hybrid 2, while the susceptible hybrid difference was 13.3. These differences inform that the susceptible hybrid is performing more than half of a magnitude worse than the tolerant hybrids when exposed to iron deficiency stress, mimicking patterns observed in field performance of the controls under stress.

To demonstrate the utility of the assay for IDC tolerance screening method for breeding and research applications, the assay was used to phenotype a population of diverse global accessions, alongside the control hybrids (Figure 4B). Under the hypothesis that the hybrid controls are representative benchmarks of farmer-available IDC tolerance, we expected that the tolerant hybrid 1 would exhibit higher SPAD and the susceptible hybrid would exhibit lower SPAD than the majority of the test accessions. As expected, the tolerant control hybrid 1 performed substantially better than the distribution mean but not better than all accessions while the susceptible control hybrid performed substantially worse than the distribution mean. Of the 330 accessions screened, 66 accessions had higher average SPAD than the tolerant hybrid, 195 accessions exhibited average SPAD values between that of the tolerant and susceptible hybrids, and 69 accessions had lower average SPAD than the susceptible hybrid. The performance of the controls relative to the distribution of test accession phenotypes supports the functionality of the assay as a field proxy in detecting genetic variation and germplasm donors for IDC tolerance.

### Removing spatial noise increases detection of genetic differences in stress tolerance

To understand how controlling for spatial variation affects the ability to detect genetic signals for IDC tolerance, the *H^2^* of field and controlled-environment phenomics approaches (Table S1) were assessed for the hybrid controls. Under the hypothesis that spatial variation in field conditions masks genetic variation for IDC tolerance, we expect that removing phenotypic data from putatively unstressed field areas would strengthen genetic signals and increase broad-sense heritability for IDC stress tolerance phenotypes. Furthermore, we would expect that the controlled-environment assay, designed to administer equal iron deficiency stress treatment to each sample, would produce high *H^2^* for IDC tolerance phenotypes due to spatial control of the stress. We found that *H^2^* for mean Q1 NDRE increased from 0.18 in the field dataset not filtered for stress plots to 0.41 after applying the neighbor-filtering method currently used in breeding programs. Moverover, the neighbor-filtering approach performed closely to a pH-stress severity filtering approach with only 0.03 difference in *H^2^* between the field filtering methods (Figure 4C). In the controlled-environment assay, *H^2.^* for SPAD increased to 0.97-0.98 across multiple iterations of the experiment on the hybrid controls (Figure 4D). From these stepwise increases observed in *H^2^* at levels of spatial control, we recognize the positive impact that accounting for spatial variation has on detection of genetic variation for IDC tolerance in sorghum.

### Controlled-environment assay facilitates testing and generation of genetic hypotheses

To determine the utility of the controlled-environment assay to inform genetic discoveries, a genome-wide association study (GWAS) and colocalization analysis were completed based on phenotype data collected from the assay. We expected that if monogenic or oligogenic trait variation for IDC tolerance was present in the georeferenced global accessions then marker-trait associations (MTA) would be detectable. Furthermore, we expected that MTAs would colocalize with *a priori* candidate genes important to iron homeostasis in grasses. We found that non-zero heritability supported distinct MTAs colocalizing with *a priori* candidates as well as loci not previously considered (Figure 4E). For example, an MTA peak is approximately 37 kb downstream of a sorghum homolog of the barley mugineic acid biosynthesis gene *Ids3* (Nakanishi et al., 1993) near the end of chromosome 8. While *Ids3* was an *a priori* candidate, an MTA peak on chromosome 7 is less than 4 kb downstream of a sulfite oxidase gene not previously considered for IDC tolerance in sorghum. Thus, controlled-environment assay facilitates hypothesis testing on genetic variation in sorghum homologs of iron homeostasis as well as generate hypotheses on additional loci potentially underlying variation for iron deficiency response.

## DISCUSSION

### Controlling for spatial variability of soil improves detection of genetic variation for IDC tolerance

We seek to inform and improve breeding efforts for IDC tolerance to secure sorghum as a viable crop on calcareous soils that currently limit production and planting in an otherwise suitable agroecological niche. Spatial variation in iron deficiency stress confounds field phenotyping and selection for IDC tolerance in sorghum, as it becomes unclear whether plants are healthy and green due to presence of adaptive genetics or simply because they are not exposed to stress (Williams et al., 1987). Drawing upon the previously characterized relationship between high pH and iron speciation to the biounavailable ferric form (Loeppert & Clarke, 1984), we characterized the spatial gradient of soil pH across a field screening nursery (Figures 2A-C) to facilitate a deeper understanding of the extent to which stress variation affects genetic signals for IDC tolerance. With soil pH as a proxy for iron deficiency stress (Hansen et al., 2003; Saleem et al., 2023), our findings show that variability in both stress severity and stress uniformity significantly reduces the power to detect genetic variation for iron deficiency chlorosis tolerance in sorghum (Figures 2F-G). This indicates that, without controlling for soil spatial variation in either the experiment itself or post-experiment data analysis, genetic variation for IDC tolerance will be masked on the premise of false positive tolerance phenotypes from areas of the field experiencing minimal or no stress.

Most sorghum breeding programs rely on testing hybrids for IDC in standard breeding nurseries across the TPE, where low check replication and lack of consistent stress pressure across years largely leaves germplasm untested for the trait. Some programs plant dedicated screening nurseries, but due to the prohibitive labor and cost requirements of grid soil sampling or proximal sensing methods (Stępień et al., 2013), these nurseries leverage highly replicated control hybrids to calibrate test genotype performance to the stress of the nearby control plots. If a susceptible control is performing equal to or better than the tolerant control in an experimental unit, test plots between them are dropped. However, it is not clearly understood if this neighbor-filtering method is effective in removing unstressed or minimally stressed plots from the model to facilitate detection of genetic variation for IDC tolerance. When evaluating the neighbor-filtering method alongside a filtering method that removes plots based on a stress severity soil pH threshold, our findings show comparable detection of genetic signals between the two approaches that both significantly improve upon those from an unfiltered field dataset (Figure 4C). However, the complex nature of the soil landscape and its effects on iron deficiency stress in plants complicates the interpretation of soil information for use in plant breeding (Lindsay, 1984).

While breeding programs may prioritize other activities and expenses over soil sampling, calibrating test genotypes to heavily replicated control plots is a suitable and efficient method to improve detection of genetic signals for IDC tolerance amidst soil spatial variation in the field (Figure 4C). However, compared to the field-based methods, further isolating the effects of iron bioavailability on test genotypes (Luna et al., 2018) using a controlled-environment assay substantially improves detection of genetics underlying IDC tolerance in sorghum (Figure 4D). This suggests that the field and lab-based approaches both contribute valuable insights to the IDC tolerance breeding pipeline for sorghum.

### Dissecting IDC tolerance via acquisition of ferric iron with a controlled-environment assay

The widespread utilization of iron in plants for molecular processes creates numerous hypotheses on the genetics underlying increased tolerance to IDC in sorghum (Connorton et al., 2017). The observation of IDC-stressed sorghum primarily on agricultural lands with high pH or calcareous soils paired with previous discoveries of poor mugineic acid production in sorghum compared to other grass crops (Kawai et al., 1988) supports a need to further characterize genetic variation in the iron acquisition pathway. To limit some of the complex agroecological interactions in the field environment (Vandermeer & Perfecto, 2017), we designed a high-throughput method to isolate the impacts of iron bioavailability on sorghum performance and facilitate evaluation of IDC tolerance donors for breeding in a spatially-controlled model. We found that minimizing inputs of exogenous iron in growth substrate and plant nutrition (Figure 3A) assisted in the isolation of genetic response to iron deficiency (Figure 4D), exemplified by consistent performance in control hybrids following their putative IDC tolerance designation assigned from field classification. This phenotypic segregation pattern of controls (Figure 4A) indicates that the assay successfully proxies field iron deficiency stress with appropriate bioavailable and biounavailable iron sources (Goos & Germain, 2001), though omission of other agroecological relationships in the field such as plant relationships with symbiotic mycorrhizae (Caris et al., 1998; Xie et al., 2019) may remove important elements of the plant iron uptake system.

**Figure 3.**
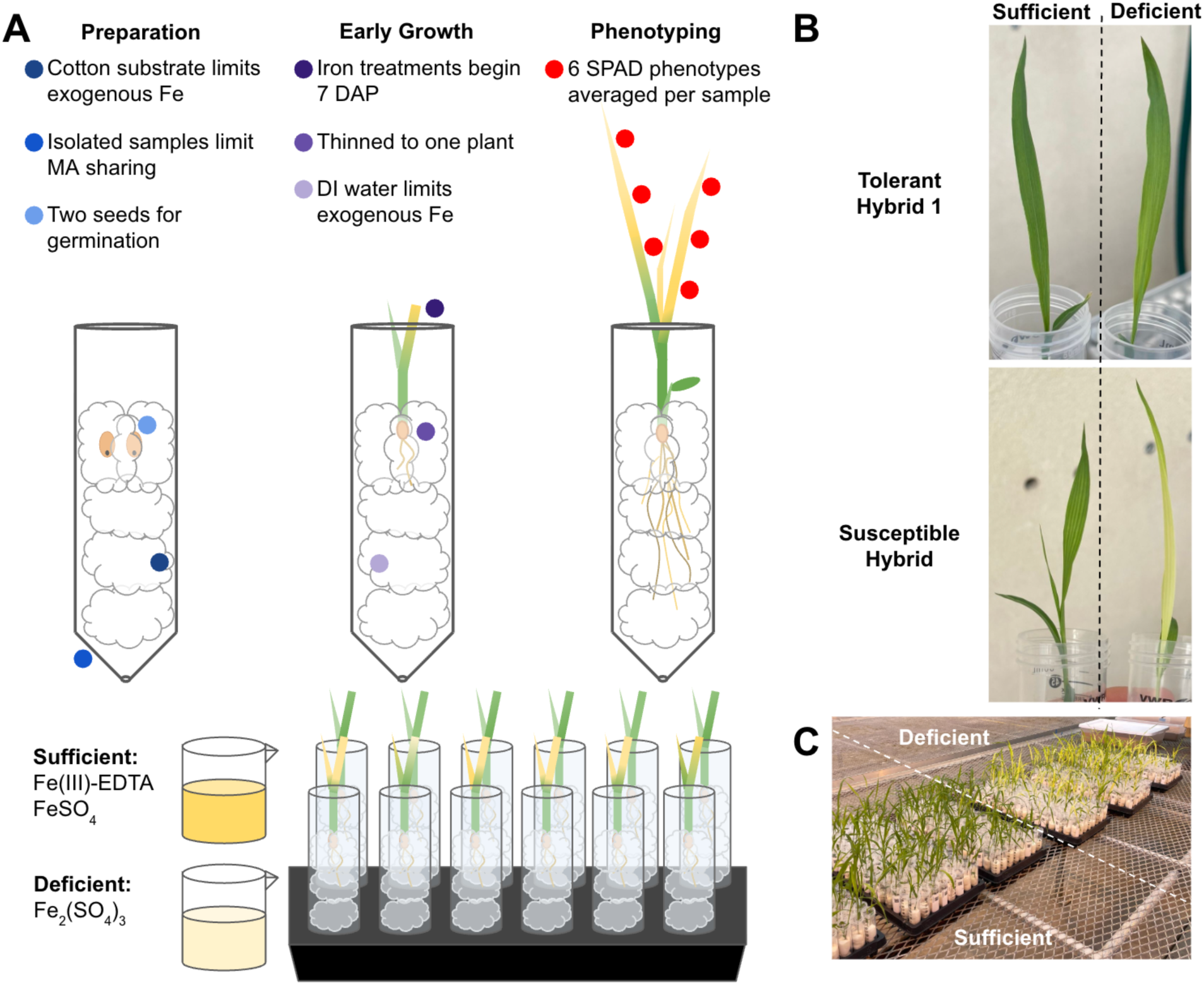
A proposed controlled-environment assay to screen sorghum germplasm for tolerance to iron deficiency chlorosis with field-relevant iron bioavailability. A) Controlled-environment assay experimental design choices aim to exclude exogenous iron from the system and isolate genotype response to iron treatments. Samples are isolated to one genotype per 50mL centrifuge tube with a hole drilled in the bottom to prevent genotypes with high mugineic acid production from rescuing genotypes with poor mugineic acid production. Samples are grown in cotton due to the primarily cellulose composition and high hydrophilicity. Application of sufficient iron treatment containing ferric-EDTA and ferrous sulfate and deficient iron treatment containing ferric sulfate begins at 7 days after planting (DAP). Deionized water (DI) water is always used for germination and nutrient solutions due to the removal of dissolved mineral ions. Given iron deficiency affects young plant tissues, phenotypes are collected approximately one month after planting where three SPAD measurements from each of the two youngest leaves are averaged for a sample phenotype. **B)** Control hybrid performance in controlled-environment assay under sufficient and deficient treatments. **C)** Screening a panel of georeferenced global sorghum accessions using the controlled-environment assay.

**Figure 4.**
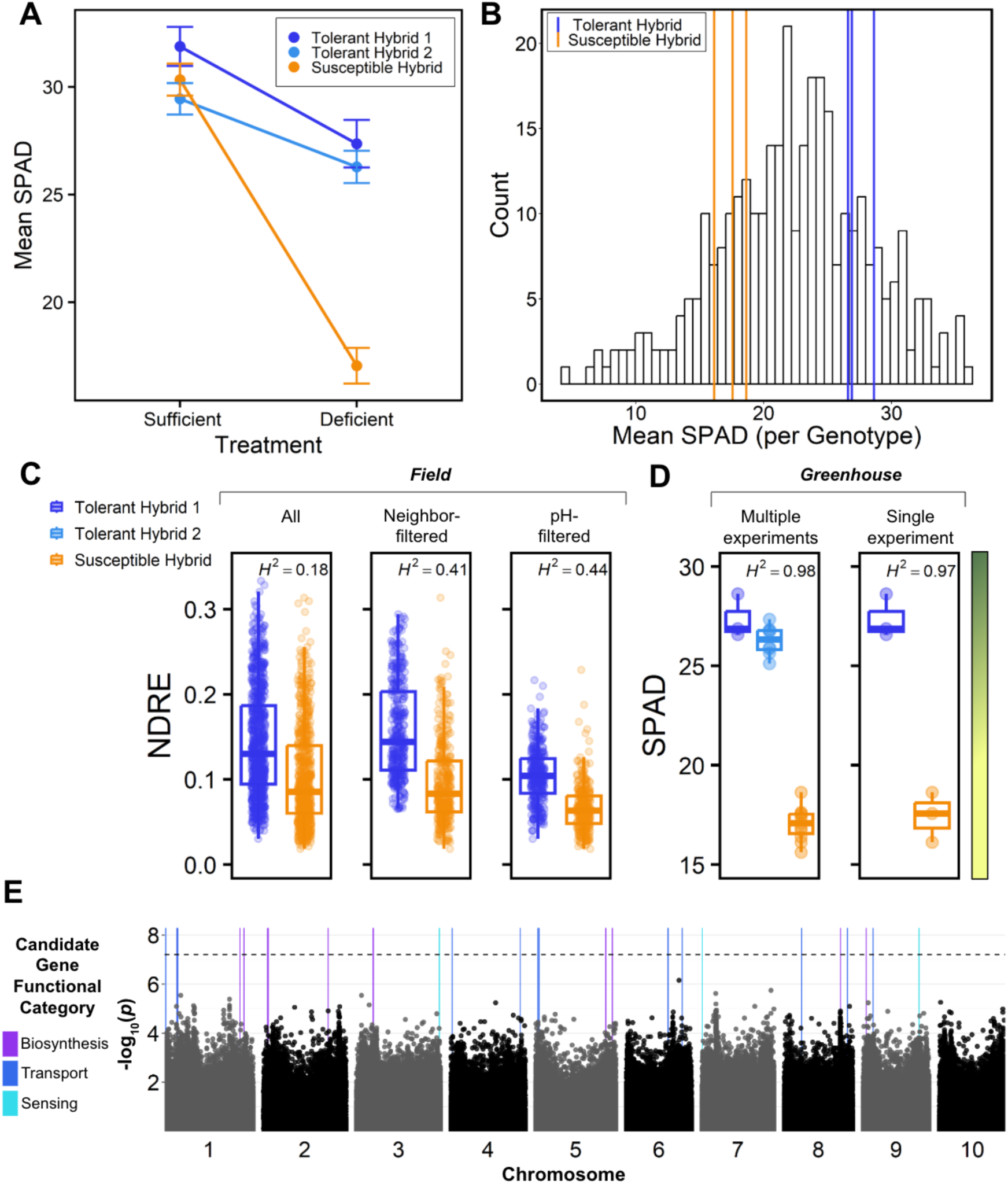
The controlled-environment assay adequately proxies field iron deficiency chlorosis stress and facilitates detection of IDC tolerance genetics across sorghum breeding material and diverse accessions. **A)** Tolerant hybrid controls significantly outperform the susceptible hybrid control under iron deficiency treatment in controlled-environment assay (per treatment: tolerant hybrid 1 n = 3, tolerant hybrid 2 n = 8, susceptible hybrid n = 11). **B)** Putative tolerance and susceptibility of control hybrids to IDC is applicable to germplasm beyond commercial breeding programs, such as a georeferenced sorghum diversity population phenotyped for IDC tolerance using the controlled-environment assay (n= 312). **C)** Removal of field plots by stress-filtering where the NDRE phenotype of the tolerant and susceptible hybrids is not significantly different or the susceptible hybrid performance is higher than the tolerant hybrid performance increases detection of genetic signals for IDC tolerance relative to no filtering. Removal of plots by stress-filtering according to a pH threshold where putatively unstressed plots are dropped also increases detection of genetic signals for IDC tolerance relative to no filtering. Tolerant hybrid 1: all n = 798, neighbor-filtered n = 358, pH-filtered n = 335. Susceptible hybrid: all n = 669, neighbor-filtered n = 358, pH-filtered n = 305. **D)** Performance of hybrid controls under assay iron deficiency treatment mimics phenotype patterns observed in managed stress field performance where tolerant controls greatly surpass susceptible control performance. Yellow-green bar is a visual representation of plant greenness across the observed SPAD scale. Tolerant hybrid 1: multiple experiments n = 3, single experiment n = 3. Tolerant hybrid 2: multiple experiments n = 8. Susceptible hybrid: multiple experiments n = 11, single experiment n = 3. **E)** Genome-wide associate study for SPAD in a panel of georeferenced global sorghum accessions (n = 312).Vertical lines represent positions of *a priori* IDC tolerance candidate genes, colored by functional role in iron homeostasis pathways.

With the understanding that the controlled-environment assay reduces agroecological complexity present in field landscapes, the differential iron assay we developed is a valuable tool for identification of candidate trait donor germplasm for breeding increased IDC tolerance in sorghum. The widespread susceptibility of sorghum to IDC in U.S. production regions suggests a lack of genetic variation for tolerance in elite breeding material (Obour et al., 2019). Therefore, our assay is not only valuable to screen existing breeding program material for performance under stress, but also to identify sources of increased IDC tolerance from prebreeding and exotic sorghum germplasm (Figure 4B). Exotic germplasm does not benefit from the heterosis conferring increased vigor to hybrids (Schnable & Springer, 2013), therefore any diverse germplasm found to outperform the tolerant hybrid control in the IDC tolerance screening assay can be treated as a viable candidate IDC tolerance trait donor. Furthermore, phenotypes collected from the controlled-environment assay can be leveraged in genome wise association analyses to characterize the genetic architecture of IDC tolerance and subsequently identify specific genetic candidates to track in breeding programs (Figure 4E).

### Conclusions

Signals for genetic variation shaping sorghum tolerance to iron deficiency chlorosis are notably obscured by spatial heterogeneity of soil characteristics in field experiments (Williams et al., 1987). The year-to-year variability of IDC stress in field environments makes reliable selection in breeding nurseries difficult and often limits progress in breeding for IDC tolerance. Controlled-environment approaches that deliberately reduce spatial noise can improve characterization of genetic responses and phenotypic performance of trait donors under iron deficiency. Our assay supports breeding efforts with multifaceted utility in evaluating existing elite germplasm for hybrid parents, screening diverse germplasm for candidate trait donors, and providing spatially-controlled phenotypes for genetic analyses. Additional characterization of field stress heterogeneity across years would further strengthen identification of trait donors in screening nurseries. Integration of tandem field and controlled-environment phenomic approaches facilitates effective breeding for increased IDC tolerance in sorghum through rapid identification of candidate tolerance sources in the lab and spatially-informed assessment of germplasm performance in on-farm field settings across the TPE (J. T. Anderson et al., 2014).

## SUPPLEMENTARY DATA

**Table S1.**
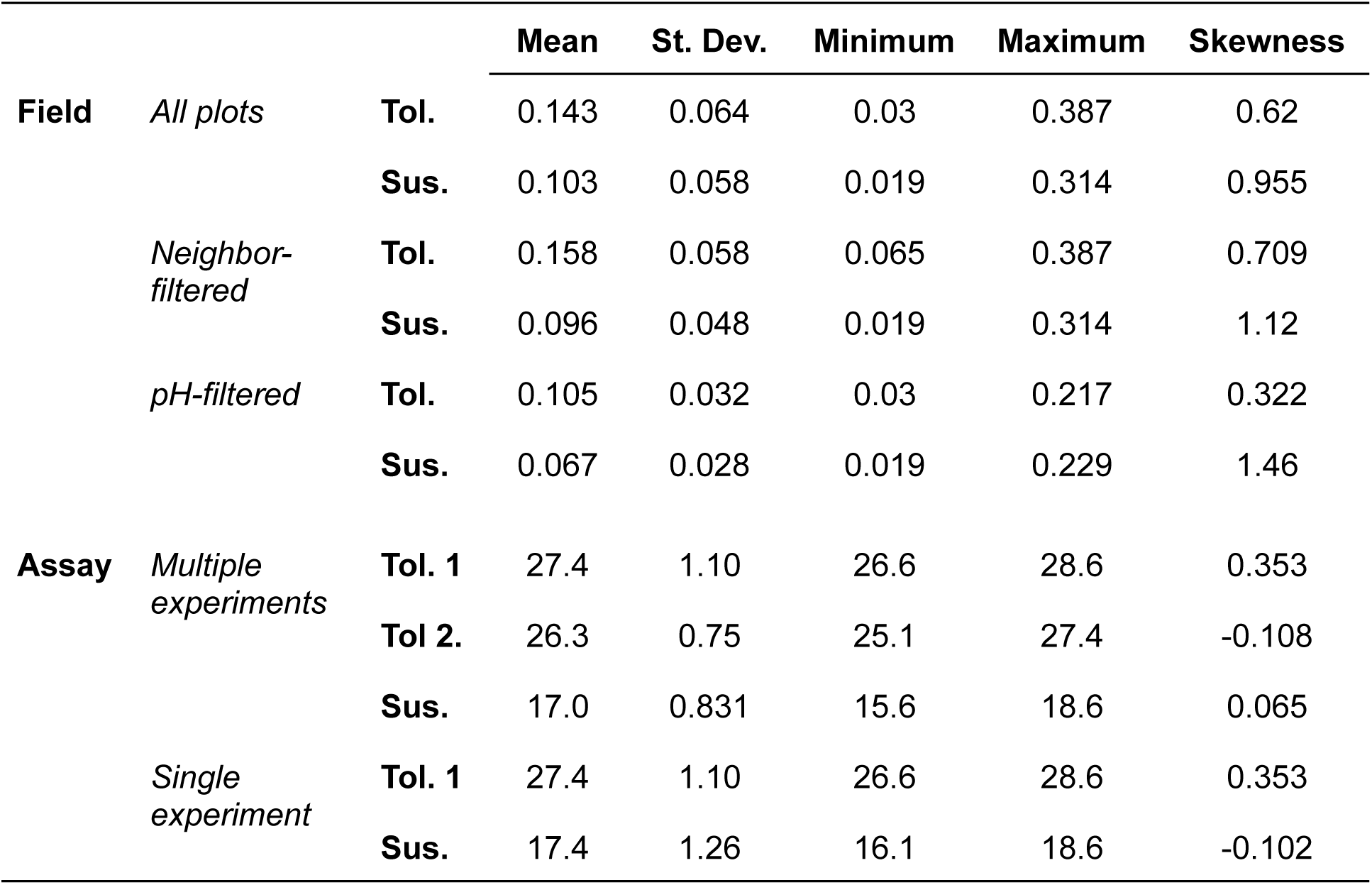
Summary statistics of phenotypes across levels of spatial control.

## ACKNOWLEDGEMENTS

We thank Dr. Lucas Haag of the Kansas State University Northwest Research-Extension Center for his management of the field site and providing the Veris soil data from the site.

## AUTHOR CONTRIBUTION

GC and GPM conceived the study. MK, MM, RB, and CF provided breeding program insights, field imagery and phenotype data, and hybrid seed. NK and TF provided extracted soil pH data. GC designed the controlled-environment assay and carried out experiments, and completed all analyses. MK, NK, TF, SS, GC, and GPM edited the manuscript.

## DATA AVAILABILITY

Data is available at [insert Dryad accession after acceptance] and analysis scripts are available at [insert GitHub link after acceptance]

